# Phage Display-Derived Cyclic Peptides Target TREM2 and Modulate Microglial Responses under Amyloid Stress

**DOI:** 10.64898/2026.04.22.720287

**Authors:** Natalie Fuchs, Shaoren Yuan, Katarzyna Kuncewicz, Mohamed Atta Elhamouly, Farida El Gaamouch, Moustafa T. Gabr

**Affiliations:** Department of Radiology, Molecular Imaging Innovations Institute (MI3), Weill Cornell Medicine, New York, NY 10065, USA.; Department of Biomedical Chemistry, Faculty of Chemistry, University of Gdansk, Poland; Biotech Drug Research Center, Shanghai Institute of Materia Medica, Chinese Academy of Sciences, Shanghai 201203, China

**Author notes:** These authors contributed equally to this work (Natalie Fuchs, Shaoren Yuan, Katarzyna Kuncewicz).

**Keywords:** TREM2, Alzheimer’s disease, peptides, phage display, microglia

## Abstract

Triggering receptor expressed on myeloid cells 2 (TREM2) is a key regulator of microglial function and a promising therapeutic target in Alzheimer’s disease. While current strategies have largely focused on antibody-based agonists, alternative modalities capable of modulating TREM2 signaling remain underexplored. Here, we report the discovery of TREM2-binding cyclic peptides using a disulfide-constrained phage display library. Screening and biophysical validation identified multiple binders, with **TREM2-6** and **TREM2-12** exhibiting micromolar affinity. Both peptides modulated microglial responses in human iPSC-derived model of amyloid stress and in neuron-microglia co-cultures. Molecular dynamics simulations supported stable peptide-TREM2 interactions, with **TREM2-12** displaying a more constrained binding mode. In vitro pharmacokinetic profiling revealed favorable plasma and intestinal stability but limited permeability, consistent with cyclic peptide scaffolds. Together, these findings establish cyclic peptides as a viable modality for targeting TREM2 and provide a foundation for the development of tunable neuroimmune therapeutics.

**Insert Table of Contents artwork here**

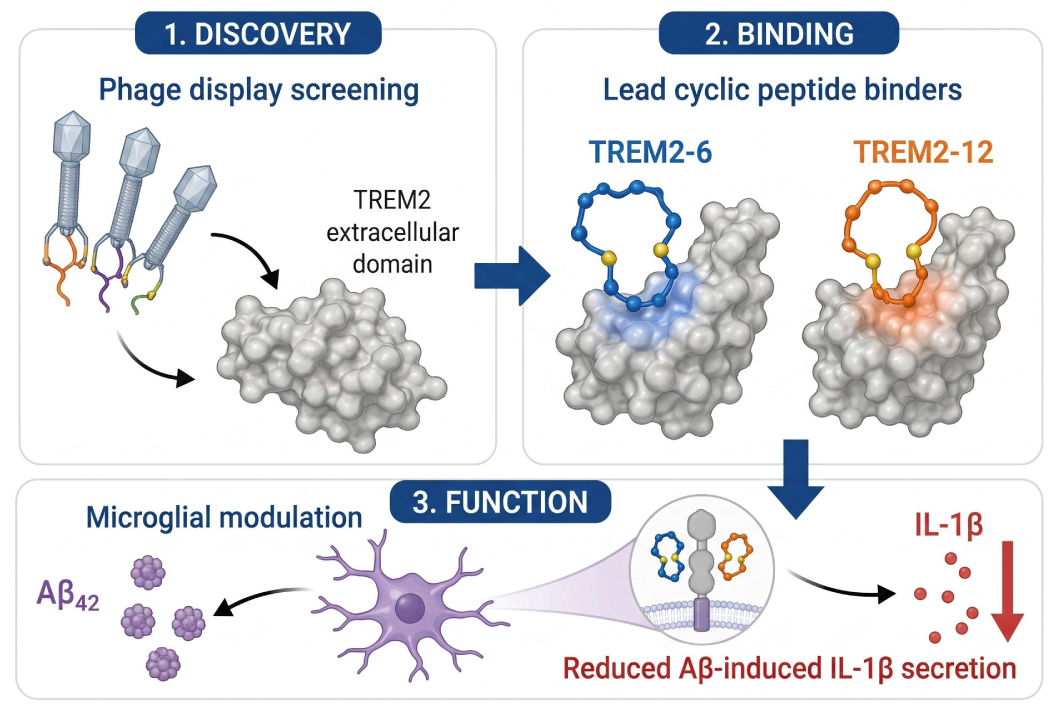

## 1. Introduction

Peptide-based therapeutics have emerged as a compelling modality bridging the gap between small molecules and biologics.^1–4^ Among these, cyclic peptides offer distinct advantages due to their conformational rigidity, which enhances proteolytic stability, improves pharmacokinetic behavior, and often increases binding affinity toward protein surfaces.^5^ Unlike linear peptides, cyclic scaffolds are particularly well suited to engage extended or shallow interfaces typical of protein-protein interactions (PPIs), enabling modulation of targets that are traditionally considered difficult to drug.^6^ Despite their promise, the discovery of functional cyclic peptides remains challenging, largely due to the vast sequence space and the need for efficient platforms that can identify binders with both high affinity and functional relevance.

TREM2 (triggering receptor expressed on myeloid cells 2) is a transmembrane receptor predominantly expressed on microglia, where it plays a central role in regulating innate immune responses in the central nervous system.^7–10^ Genetic studies have identified TREM2 variants as significant risk factors for neurodegenerative diseases, including Alzheimer’s disease, highlighting its importance in disease progression.^11,12^ Mechanistically, TREM2 signaling has been linked to microglial activation, phagocytosis, lipid sensing, and modulation of neuroinflammation.^13,14^ As a result, TREM2 has emerged as an attractive therapeutic target, with efforts largely focused on antibody-based agonists designed to enhance its signaling activity.^15^ However, biologics face limitations including restricted tissue penetration, prolonged receptor engagement, and limited control over temporal signaling dynamics.

Phage display provides a powerful platform for the discovery of peptide ligands, enabling the rapid screening of large and diverse libraries against target proteins.^16–19^ In particular, disulfide-constrained cyclic peptide libraries allow for the identification of conformationally restricted binders that can mimic structural features of protein interaction interfaces.^17,18^ These libraries have been successfully used to generate peptides with high affinity and specificity across a range of challenging targets. However, the application of phage display to microglial receptors such as TREM2 remains underexplored.

Building on our prior work targeting TREM2 with small molecules, which demonstrated robust activity across biochemical and cellular assays,^20–26^ we sought to expand into alternative modalities capable of more effectively engaging the receptor surface. In parallel, we previously reported an AI-guided cyclic peptide binder targeting TREM2;^27^ however, while this approach provided an important proof-of-concept, the resulting peptides exhibited only modest affinity in the sub-millimolar range, highlighting the challenges of rational peptide design for this target.

Here, we report the identification of cyclic peptide ligands targeting TREM2 using a phage display approach. By screening a disulfide-constrained peptide library, we identified candidates with strong binding affinity and functional activity. Lead peptides were further characterized using biophysical and cellular assays to assess their ability to modulate TREM2 signaling. This work establishes cyclic peptides as a viable alternative modality for targeting TREM2 and provides a foundation for developing tunable immunomodulators with improved control over microglial activation.

## 2. Results and discussion

### Phage display reveals cyclic peptide binders targeting the TREM2 extracellular domain

To engage the relatively featureless extracellular surface of TREM2, we performed phage display selection using a disulfide-constrained cyclic peptide library based on the CX_9_C scaffold, designed to present conformationally restricted interaction motifs compatible with extended protein-protein interfaces. The library was subjected to four rounds of biopanning against the recombinant human TREM2 extracellular domain, resulting in progressive enrichment of phage populations across successive rounds (Figure 1A). Increased phage recovery relative to control selections is consistent with selective amplification of TREM2-binding clones rather than nonspecific retention.

**Figure 1.**
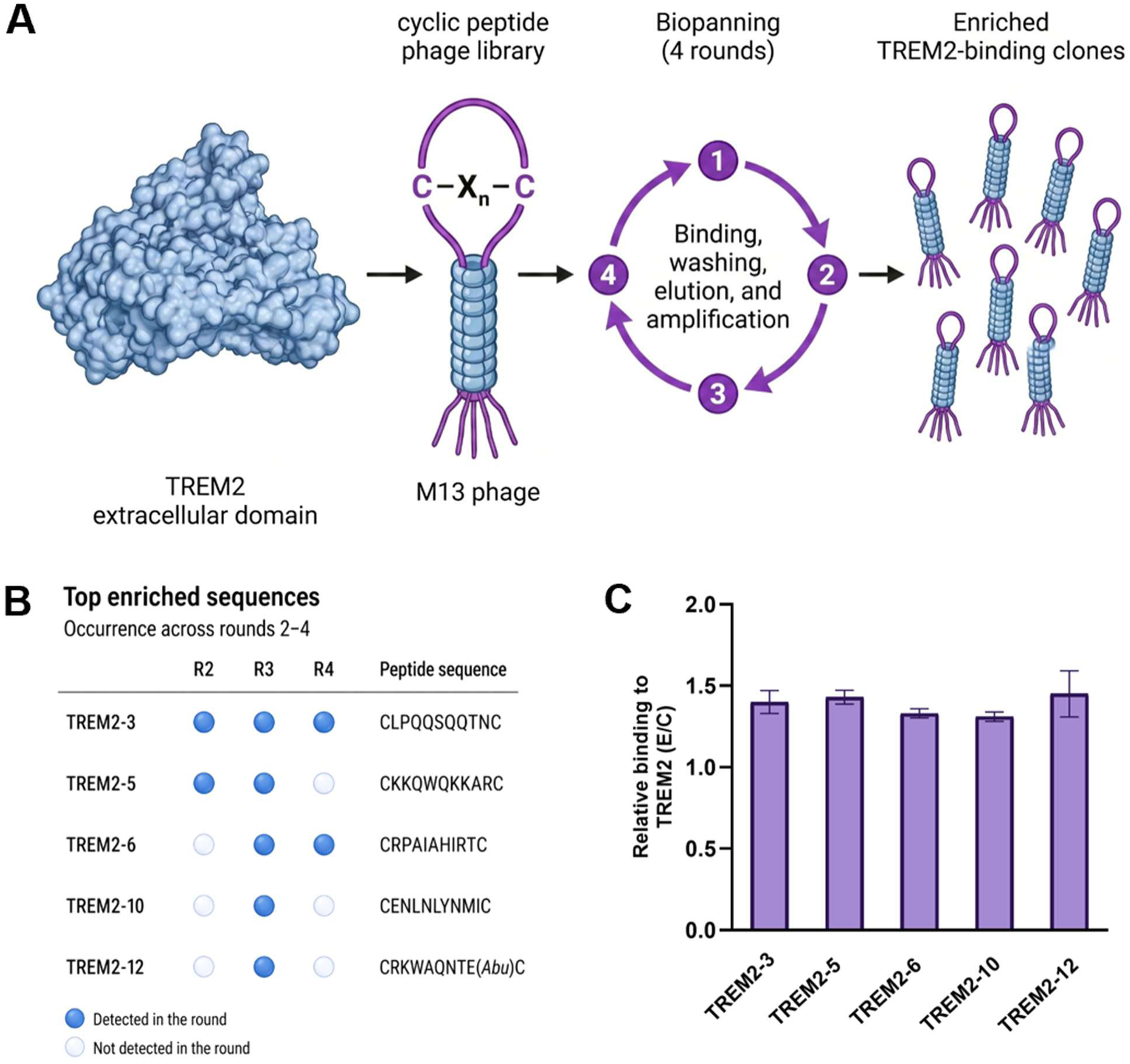
Identification and validation of TREM2-binding cyclic peptides by phage display. **(A)** Schematic of the phage display workflow. The extracellular domain of TREM2 was used as the target for selection from a cysteine-constrained cyclic peptide M13 phage library. Four rounds of biopanning, including binding, washing, elution, and amplification, were performed to enrich TREM2-binding clones. **(B)** Top enriched peptide sequences identified from rounds 2-4. Filled circles indicate detection of a given sequence in the corresponding round, while open circles indicate absence. All sequences conform to a cysteine-constrained cyclic peptide scaffold. **(C)** Phage ELISA validation of selected peptides. Binding signals are presented as the ratio of signal obtained in TREM2-coated wells relative to control wells lacking protein (E/C). Values above 1 indicate preferential binding to TREM2. Data are shown as mean ± SEM (n=3).

Beginning in round two, individual clones were isolated and sequenced, revealing a diverse set of cyclic peptide sequences detected across subsequent rounds of selection. From this pool, a subset of representative five peptides (**TREM2-3**, **TREM2-5**, **TREM2-6**, **TREM2-10**, and **TREM2-12**) was selected for further analysis based on recurrence across rounds and sequence diversity (Figure 1B). Several of these peptides were identified in multiple independent rounds, indicating reproducible enrichment of TREM2-binding candidates despite the absence of convergence to a single dominant motif. All selected sequences retained the cysteine-constrained architecture while exhibiting substantial diversity, suggesting that TREM2 engagement can arise from multiple structurally distinct solutions rather than a single privileged binding mode.

Functional interrogation by phage ELISA confirmed that all selected peptides exhibited measurable binding to immobilized TREM2 (Figure 1C). Binding signals, expressed as the ratio of signal obtained in TREM2-coated wells relative to control wells lacking protein (E/C), were consistently above baseline, although magnitudes varied across peptides. This indicates that sequence enrichment alone does not directly predict binding strength and highlights the presence of a functionally selective subset of ligands. Collectively, these results demonstrate that cyclic peptide phage display can identify experimentally validated binders to the TREM2 extracellular domain and enable prioritization of candidates for downstream biophysical and functional characterization.

### Monolith assay for TREM2 binding

We initially evaluated a prioritized set of ten peptides derived from the phage display output, comprising the five most enriched sequences shown in Figure 1B and five additional candidates selected to represent sequence diversity and avoid bias toward a single enrichment trajectory. This initial evaluation was performed in a single-dose screening at 50 µM using Monolith X (raw data in Figure S1, Supporting Information). The assay buffer (HBSN pH 7.4, n = 4) served as a negative control whereas previously reported TREM2 binder **T2337** (50 µM, n = 2) was used as a positive control.^24^ Peptides with a spectral shift (SpS, ratio 670 nm/650 nm) outside of a five standard deviation range from the average negative control were considered as potential hits. In the screening, six peptides were outside of the range, but **TREM2-11** was flagged by the software for aggregation and adsorption to the capillaries, resulting in its exclusion from further experiments (Figure 2A). Furthermore, all hit peptides caused a red or bathochromic shift (Figure 2B), which correlates with an increase in the local hydrophobicity of the dye, possibly due to the displacement of the hydration water shell around the dye upon peptide binding to TREM2.

**Figure 2.**
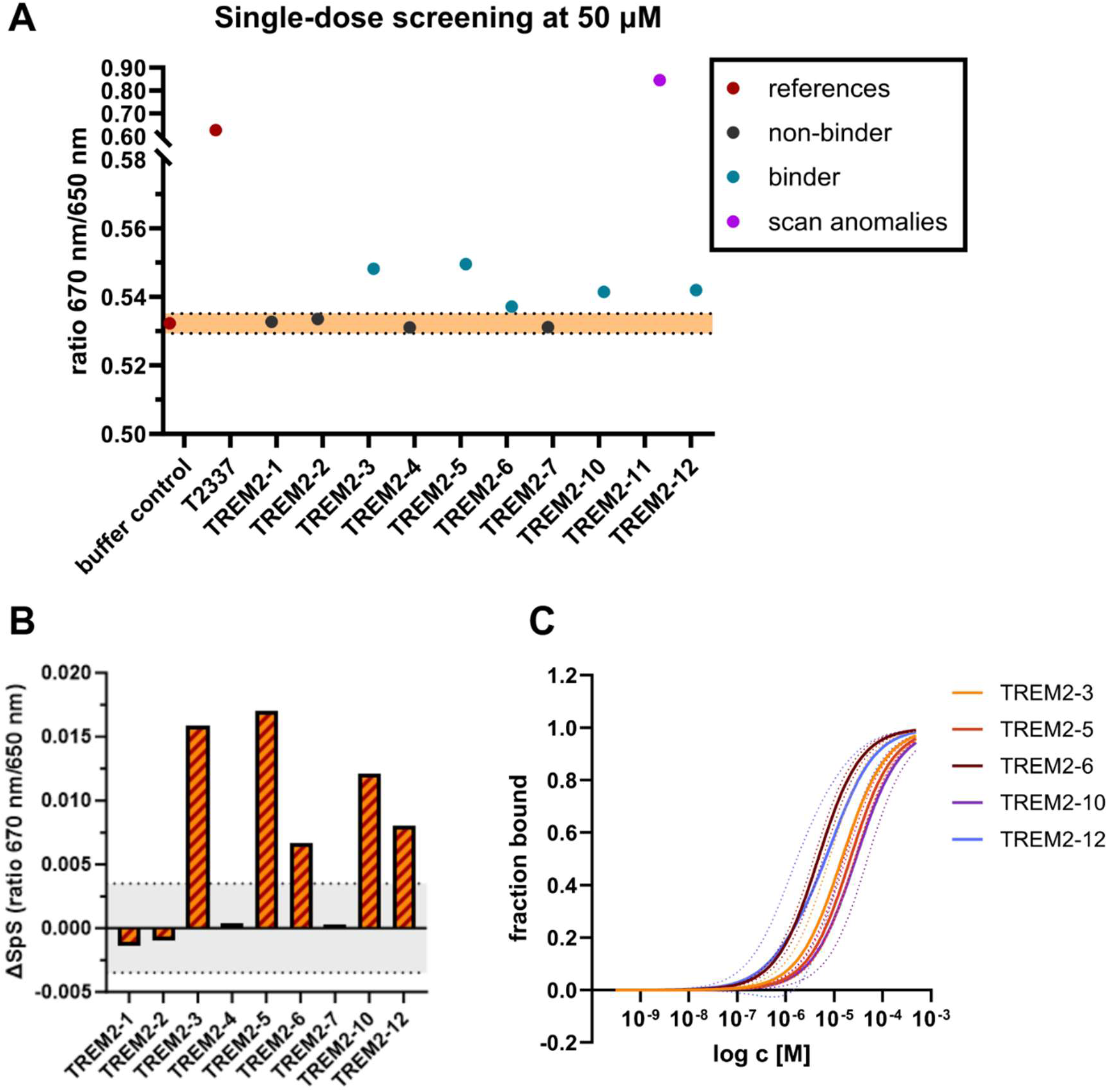
Results for Monolith binding studies. (**A**) Single-dose screening at 50 µM. Five standard deviation range from average negative control is displayed in orange with dotted lines at the upper and lower limits, potential binders as green dots, peptides that showed scan anomalies in purple. Raw data (SpS and TRIC traces) are shown in the Supporting Information (Figure S1). (**B**) Spectral shift difference (ΔSpS) from average negative control in the single-dose screening. Five standard deviation range is highlighted in gray. (**C**) Normalized fraction bound vs. log c [M] graphs for hit peptides after dose-dependent studies. Individual SpS graphs in Supporting Information (Figure S3). All graphs were created with GraphPad Prism 10. Figure composed using BioRender.

In the next step, we tested the remaining five peptides (**TREM2-3**, **TREM2-5**, **TREM2-6**, **TREM2-10**, and **TREM2-12**) in control experiments where they did not show any assay interference (see Figure S2, Supporting Information). Importantly, the peptides advanced from MST screening correspond to the top enriched sequences identified by phage display (Figure 1B), providing orthogonal validation of the selection outcome. Thus, we considered these five peptides for dose-dependent binding affinity studies. For dose-dependent studies, we prepared a 16-point dilution series starting at 100 µM for each peptide. All peptides have K_D_ values in the micromolar range with **TREM2-6** (K_D_ = 16.4 ± 4.89 µM) and **TREM2-12** (10.2 ± 1.24 µM) emerging as the top peptides in terms of TREM2 binding potency (Table 1, Figure 2C). The individual TREM2 binding curves for the five peptides are displayed in Figure S3, Supporting Information.

**Table 1.** TREM2 binding affinity overview for hit peptides using Monolith X.

| Peptide | K<sub>D</sub> (SpS) [μM]<sup>1</sup> |
| --- | --- |
| **TREM2-3** | 48.3 ± 20.1 |
| **TREM2-5** | 41.7 ± 10.5 |
| **TREM2-6** | 16.4 ± 4.89 |
| **TREM2-10** | 57.8 ± 13.9 |
| **TREM2-12** | 10.2 ± 1.24 |

<sup>1</sup>Data are shown as mean ± SEM (n=3).

### Functional evaluation of TREM2-targeting cyclic peptides

We next evaluated the biological activity of the two most promising peptides, **TREM2-6** and **TREM2-12**, to determine whether their binding interactions with TREM2 translated into functional modulation in a disease-relevant context. Using a human iPSC-derived microglial model of amyloid stress, cells were challenged with Aβ_1-42_ oligomers in the presence of increasing concentrations of each peptide, and inflammatory output was assessed by quantifying IL-1β secretion.

Consistent with the behavior observed for a clinical small molecule TREM2 agonist (VG-3927), both **TREM2-6** and **TREM2-12** produced a measurable attenuation of Aβ-induced IL-1β release in a dose-dependent manner relative to vehicle-treated controls (Figure 3A). The magnitude and dose dependence of this effect indicate that engagement of TREM2 by these cyclic peptides is sufficient to modulate microglial inflammatory responses under amyloid stress conditions. While the overall activity profiles of the two peptides were broadly similar, **TREM2-12** consistently exhibited a modestly stronger effect across the tested concentration range (5-25 μM), whereas **TREM2-6** retained clear but slightly reduced activity. Importantly, these differences were incremental rather than qualitative, suggesting that both peptides operate within a comparable functional regime.

**Figure 3.**
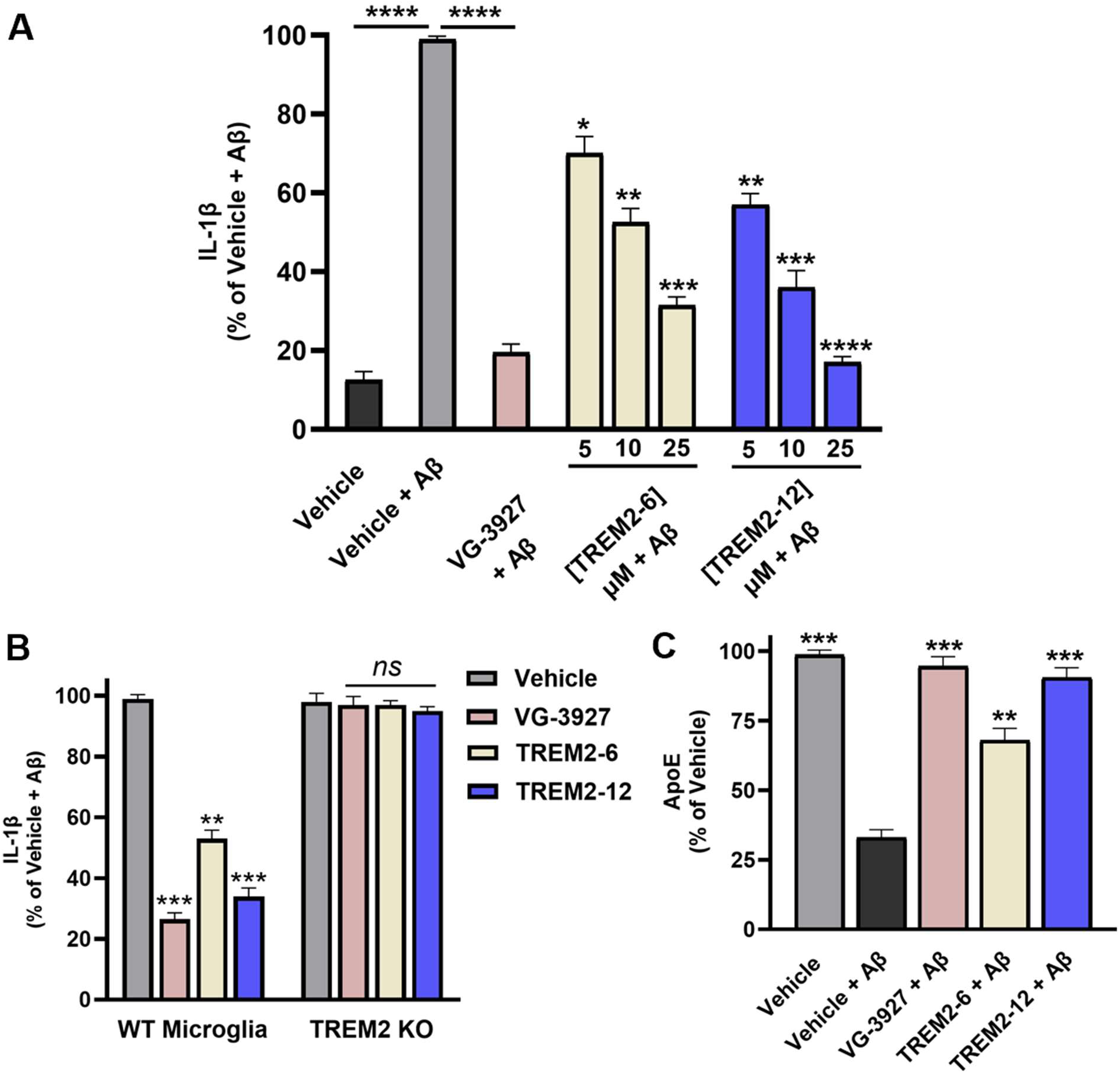
Cyclic peptides modulate Aβ-induced microglial responses in a TREM2-dependent manner. **(A)** IL-1β secretion in human iPSC-derived microglia following Aβ_1-42_ oligomer challenge in the presence of **TREM2-6** or **TREM2-12** (5, 10, and 25 µM). VG-3927 (5 μM) was used as a positive control. Data are normalized to the Aβ-treated vehicle control and expressed as percent change. **(B)** Loss of peptide-mediated suppression of IL-1β secretion in TREM2 knockout (KO) microglia confirms TREM2-dependent activity. **(C)** Modulation of secreted ApoE levels in human microglia following Aβ exposure and peptide treatment (25 µM). ApoE levels are normalized to vehicle-treated controls. Data are presented as mean ± SD (n = 5). Statistical significance was determined by one-way or two-way ANOVA with Dunnett’s post hoc test. *ns*, not significant; *p* < 0.05 (*), *p* < 0.01 (**), *p* < 0.001 (***), and *p* < 0.0001 (****) relative to vehicle treatment.

To confirm that the observed anti-inflammatory effects were mediated through TREM2, we evaluated peptide activity in TREM2 knockout (KO) microglia. Under Aβ challenge, both peptides significantly reduced IL-1β secretion in wild-type cells; however, this effect was markedly attenuated in TREM2 KO cells (Figure 3B), providing direct genetic evidence that the observed activity is TREM2-dependent rather than arising from nonspecific anti-inflammatory effects.

Given the established role of the TREM2-ApoE axis in regulating microglial activation states, we next assessed whether peptide treatment modulated ApoE secretion. Both **TREM2-6** and **TREM2-12** altered ApoE levels in Aβ-challenged microglia, with **TREM2-12** again showing a modestly stronger effect (Figure 3C). These findings indicate that peptide-mediated TREM2 engagement influences not only inflammatory cytokine output but also key lipid-associated pathways implicated in disease-associated microglial phenotypes.

We next examined whether these effects translated into functional protection in a more complex cellular system. In human iPSC-derived neuron-microglia co-cultures, Aβ exposure resulted in a pronounced reduction in PSD95 levels, consistent with synaptic damage. Treatment with both peptides led to a significant preservation of PSD95, with **TREM2-12** again showing a modestly improved effect (Figure 4). These results suggest that modulation of microglial activity by TREM2-targeting peptides can confer downstream neuroprotective benefits. Taken together, these data demonstrate that **TREM2-6** and **TREM2-12** are not only biophysical binders but also functionally active modulators of microglial behavior across multiple complementary readouts. The convergence of anti-inflammatory effects, TREM2-dependent activity, modulation of ApoE signaling, and preservation of synaptic markers provides strong evidence that cyclic peptide engagement of TREM2 can influence disease-relevant pathways. While **TREM2-12** consistently exhibited modest improvements across assays, the overall similarity in performance suggests that further optimization will likely involve incremental refinement rather than large stepwise gains. Nonetheless, both peptides establish a robust foundation for the development of next-generation TREM2-targeting agents and support the broader feasibility of cyclic peptides as modulators of neuroimmune signaling.

**Figure 4.**
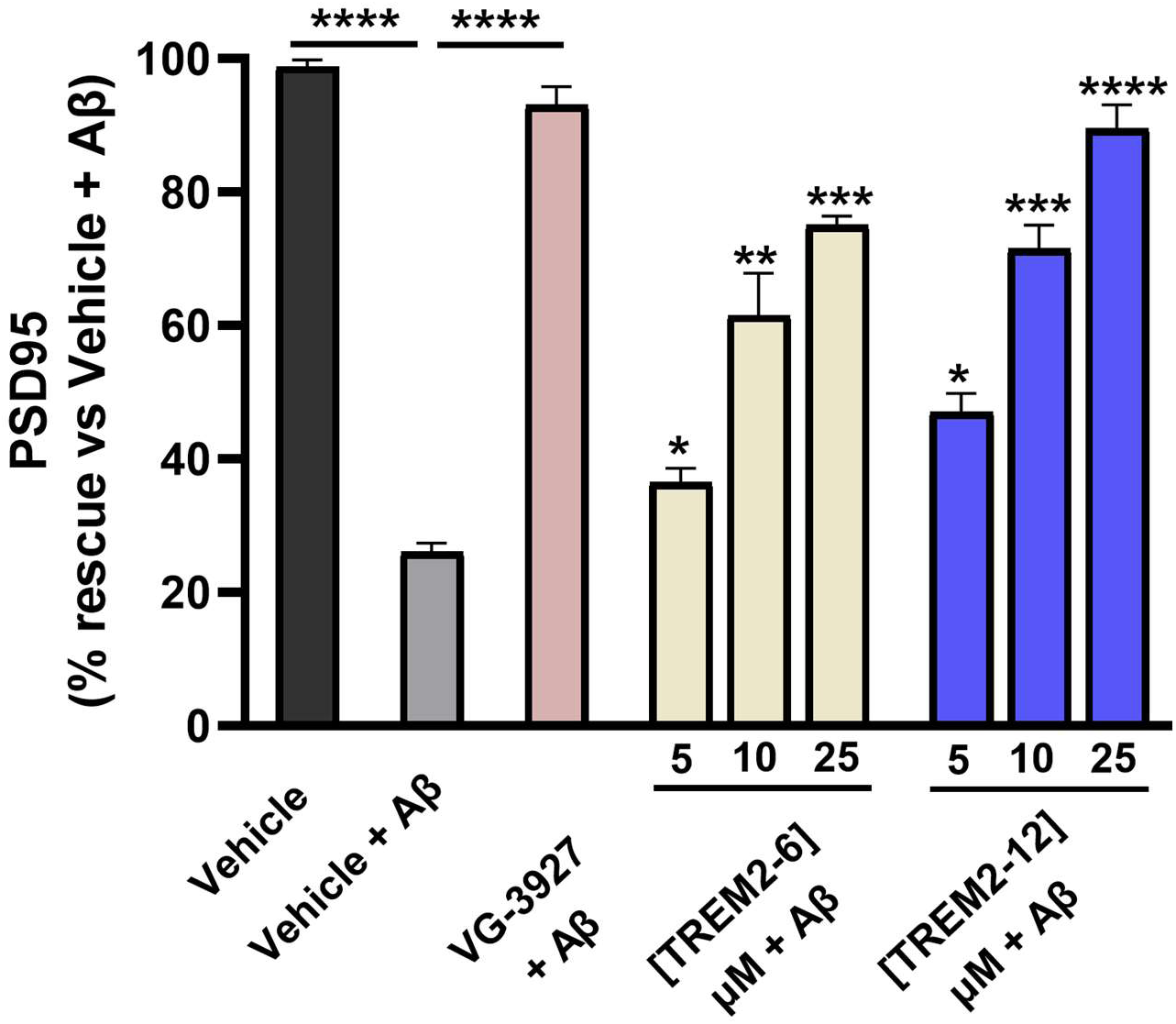
Cyclic peptides preserve synaptic markers in neuron-microglia co-cultures under amyloid stress. Quantification of PSD95 levels in human iPSC-derived neuron-microglia co-cultures following Aβ_1-42_ oligomer exposure and treatment with **TREM2-6** or **TREM2-12** (5, 10, and 25 µM). VG-3927 (5 μM) was used as a positive control. PSD95 levels are expressed as percent rescue relative to Aβ-treated vehicle controls. Data are presented as mean ± SD (n = 5). Statistical significance was assessed by one-way ANOVA with Dunnett’s post hoc test. *p* < 0.05 (*), *p* < 0.01 (**), *p* < 0.001 (***), and *p* < 0.0001 (****) relative to vehicle treatment.

### Computational analysis of TREM2-peptide interactions

To gain structural insight into cyclic peptide binding to TREM2, we performed molecular dynamics (MD) simulations on AlphaFold3-predicted complexes of TREM2 bound to **TREM2-6** (**T6**) and **TREM2-12** (**T12**) (Figure 5). Simulations were conducted for 100 ns under physiological conditions to assess the stability and dynamic behavior of each complex. Analysis of the backbone RMSD of the TREM2 receptor revealed that the apo system remained stable, with a mean RMSD of 1.52 Å (Figure 5A). Upon peptide binding, both complexes exhibited reduced structural fluctuations, with mean RMSD values of 1.33 Å for TREM2–**T6** and 1.29 Å for TREM2–**T12**, indicating that both peptides contribute to stabilization of the TREM2 extracellular domain.

**Figure 5.**
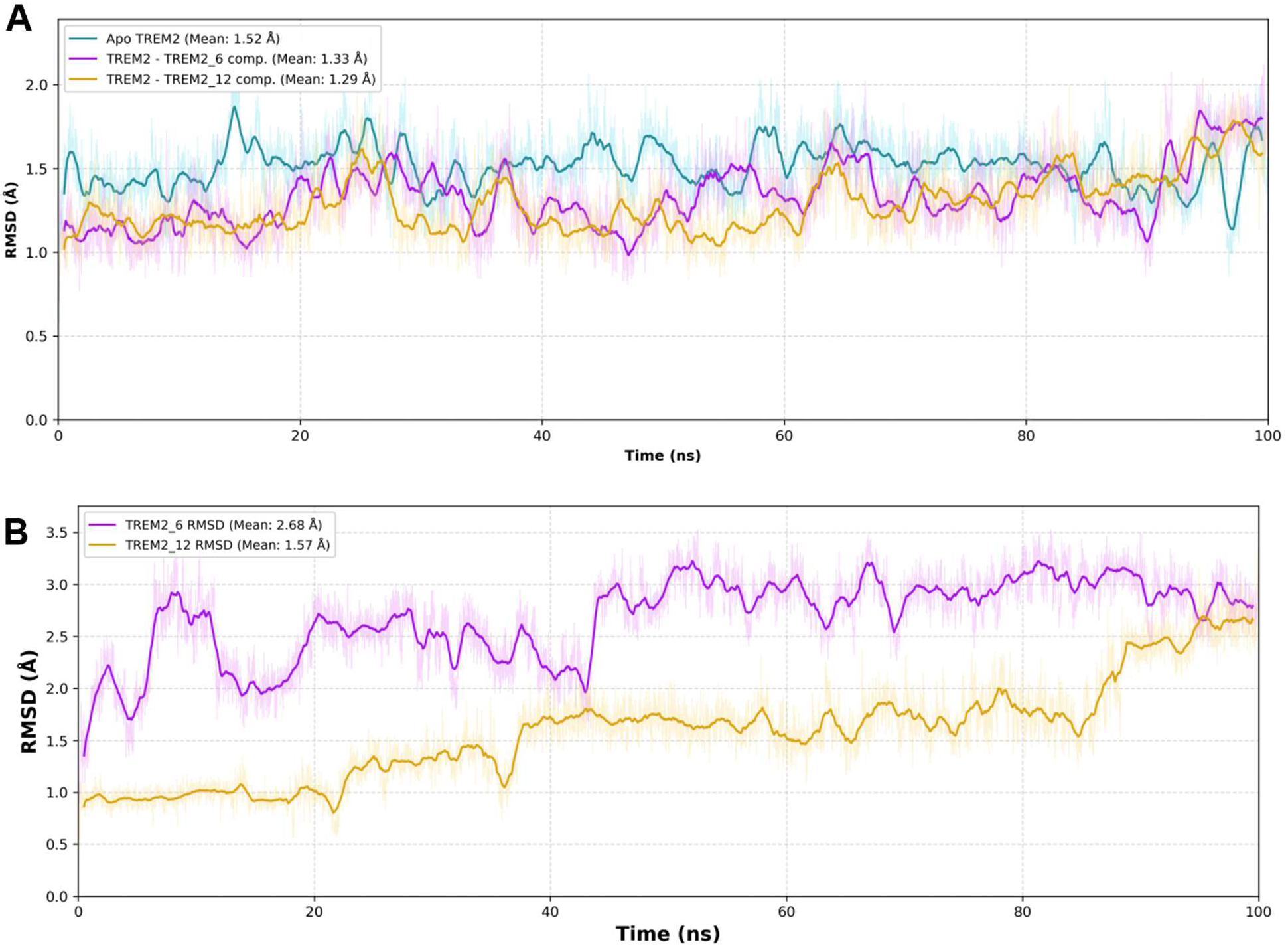
Complex prediction and MD simulation of TREM2 in complex with T6 and T12. **(A)** Time-dependent root-mean-square deviation (RMSD) of the TREM2 receptor in the free form (cyan line) and in complex with **T6** (purple line) and **T12** (yellow line) over a 100 ns MD simulation. **(B)** RMSD of the **T6** and **T12** over a 100-ns MD simulation.

Comparison of ligand dynamics showed differences in conformational behavior between the two peptides (Figure 5B). **TREM2-12** (**T12**) maintained a relatively stable binding conformation throughout the simulation, with a mean heavy-atom RMSD of 1.57 Å, whereas **TREM2-6** (**T6**) displayed increased conformational variability, with a mean RMSD of 2.68 Å. This suggests that **TREM2-12** potentially adopts a more conformationally constrained binding mode, while **TREM2-6** samples a broader conformational space within the binding interface.

Representative structures extracted from equilibrated trajectories (Figure 6A-D) show that both peptides engage solvent-exposed regions of the TREM2 extracellular domain through distributed interaction networks. **TREM2-12** forms multiple hydrogen bonds with residues including Arg44, Cys42, Gln93, Ser98, and Glu99 (Figure 6C), whereas **TREM2-6** interacts with residues such as Arg34, Asp86, Arg80, and Arg44 (Figure 6D). These interactions indicate that both peptides bind through multi-point surface contacts rather than a single dominant interaction site, consistent with the absence of a well-defined binding pocket on TREM2. Taken together, these results indicate that both **TREM2-6** and **TREM2-12** form stable complexes with the TREM2 extracellular domain, with peptide binding contributing to global receptor stabilization. The reduced conformational variability observed for **TREM2-12** suggests a more pre-organized binding mode, which is consistent with its modestly improved binding affinity relative to **TREM2-6** observed experimentally. At the same time, the comparable overall behavior of the two peptides indicates that they operate within a similar binding regime, rather than representing fundamentally distinct interaction modes. More broadly, the ability of both peptides to engage TREM2 through distributed surface interactions supports a model in which cyclic peptides can access shallow or featureless protein interfaces through multiple sequence-dependent solutions. This is consistent with the sequence diversity observed in the phage display selection and highlights the potential for further optimization through iterative refinement rather than reliance on a single privileged motif.

**Figure 6.**
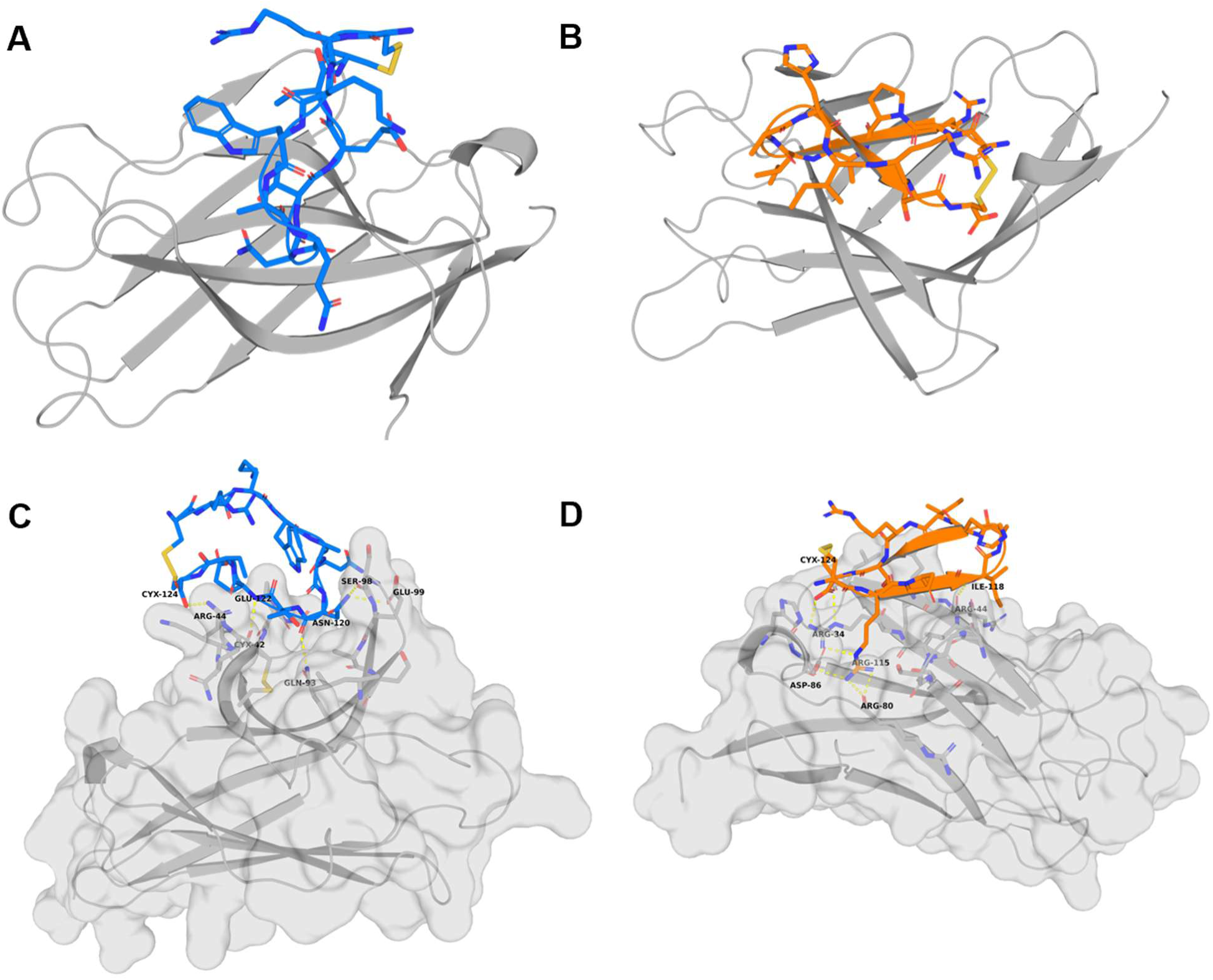
Structural characterization of TREM2-peptide interactions by MD simulations. (A and. **B)** Structural snapshot clustered from the late stage of the MD simulation of **T12 (A)** and **T6 (B)** complexed with TREM2. The receptor is shown in grey cartoon, **T12** is shown in blue sticks, and T6 is shown in orange sticks. **(C and D)** Views of the binding interface between **T12 (C)** / **T6 (D)** and TREM2 with the key interacting residues on both sides and yellow dashed lines that demonstrate stable hydrogen bonds and salt bridges.

### Pharmacokinetic profiling of the top TREM2 peptides

To gain preliminary insight into the developability of the lead cyclic peptides, we next evaluated the in vitro pharmacokinetic (PK) profiles of **TREM2-6** and **TREM2-12** (Table 2). Such analyses are important for assessing the translational potential of peptide scaffolds, as stability in biological environments, physicochemical properties, and permeability collectively influence their suitability as therapeutic agents.

**Table 2.** *In vitro* PK profile of **TREM2-6** and **TREM2-12**.

| Test | TREM2-6 | TREM2-12 |
| --- | --- | --- |
| Stability in simulated fluids ($t_{1/2}$, h) | | |
| Gastric (pH 1.6) | 0.78 | 0.89 |
| Intestinal (pH 6.5) | 4.90 | 5.92 |
| Stability in human plasma (% remaining at 1 h) | 74.6 | 79.5 |
| Aqueous solubility at pH 7.4 (μM) | 131.5 | 112.7 |
| PPB (% bound, fu) | 18.7 | 24.9 |
| Log$D_{7.4}$<sup>a</sup> | 0.84 | 1.12 |
| CACO cell permeability<sup>b</sup> |  |  |
| $P_{app, A \rightarrow B}$ ($10^{-6}$ cm/s) | 0.92 | 0.68 |
| $P_{app, B \rightarrow A}$ ($10^{-6}$ cm/s) | 1.88 | 1.72 |
| Stability in rat liver microsomes ($t_{1/2}$, h) | 2.35 | 3.14 |

<sup>a</sup>Determined by miniaturized shake-flask procedure at pH7.4.
<sup>b</sup>Permeation through a Caco-2 cell monolayer was assessed in the absorptive apical to basolateral (a→b) and secretory basolateral to apical (b→a).

Both peptides exhibited limited stability under simulated gastric conditions (t₁/₂ = 0.78 h for **TREM2-6** and 0.89 h for **TREM2-12**) but showed improved stability in simulated intestinal fluid (t₁/₂ = 4.90 h and 5.92 h, respectively), consistent with the protective effect of backbone cyclization under near-neutral pH conditions. In human plasma, both peptides demonstrated good resistance to proteolytic degradation, with 74.6% (**TREM2-6**) and 79.5% (**TREM2-12**) remaining after 1 h.

The peptides displayed satisfactory aqueous solubility at physiological pH (131.5 μM for **TREM2-6** and 112.7 μM for **TREM2-12**) and relatively low plasma protein binding (18.7% and 24.9% bound, respectively), suggesting a reasonable balance between solubility and protein association. LogD_7.4_ values of 0.84 and 1.12 for **TREM2-6** and **TREM2-12**, respectively, indicate a slightly higher lipophilicity for **TREM2-12** (Table 2).

Consistent with their cyclic peptide nature, both peptides exhibited low permeability across Caco-2 cell monolayers, with Papp (A→B) values of 0.92 × 10^-6^ cm/s for **TREM2-6** and 0.68 × 10^-6^ cm/s for **TREM2-12**. Higher transport in the basolateral-to-apical direction (Papp B→A = 1.88 and 1.72 × 10^-6^ cm/s, respectively) suggests moderate efflux, which is typical for peptide scaffolds.

Both peptides also demonstrated moderate metabolic stability in rat liver microsomes, with half-lives of 2.35 h for **TREM2-6** and 3.14 h for **TREM2-12** (Table 2). Notably, **TREM2-12** consistently showed modest improvements across several parameters, including enhanced intestinal and microsomal stability and slightly increased lipophilicity, which may be consistent with its more constrained conformation observed in MD simulations and its modestly improved binding affinity. However, these differences were incremental, and both peptides remain within a similar overall pharmacokinetic and physicochemical space.

Collectively, these results indicate that **TREM2-6** and **TREM2-12** combine favorable plasma and intestinal stability with moderate metabolic stability and limited permeability, supporting their potential as tractable cyclic peptide scaffolds for further optimization.

In this study, we report the discovery and characterization of cyclic peptide ligands targeting the TREM2 extracellular domain using a phage display approach. Screening of a disulfide-constrained peptide library enabled the identification of multiple binders, with **TREM2-6** and **TREM2-12** emerging as the most promising candidates based on biophysical and functional evaluation. Both peptides exhibited micromolar binding affinity and demonstrated the ability to modulate microglial responses under amyloid stress, reducing Aβ-induced inflammatory signaling in a human iPSC-derived model. Computational analysis supported stable peptide-receptor interactions and suggested that TREM2 engagement occurs through distributed surface contacts rather than a single dominant binding site, consistent with the diversity observed in the phage display output. In vitro PK profiling revealed favorable plasma and intestinal stability, moderate metabolic stability, and limited permeability, in line with expectations for cyclic peptide scaffolds.

Collectively, these findings establish cyclic peptides as a viable modality for targeting TREM2 and highlight their potential as tunable neuroimmune modulators. While further optimization will be required to improve affinity and PK properties, the identified peptides provide a strong foundation for the development of next-generation therapeutics targeting microglial signaling in neurodegenerative disease.

## 3. Materials and methods

### Phage display selection of TREM2-binding cyclic peptides

A disulfide-constrained phage-displayed cyclic peptide library based on a CX_9_C scaffold was used to identify ligands capable of engaging the extracellular domain of human TREM2. Recombinant human TREM2 extracellular domain (SinoBiological, 11084-H08H) was immobilized on a solid support, and the phage library was subjected to iterative rounds of biopanning against the target protein. After each round, nonbinding phages were removed by washing, whereas bound phages were recovered by elution and amplified in *E. coli* for use in the subsequent round of selection. A total of four rounds of biopanning were performed, with enrichment of TREM2-binding phage monitored across rounds.

Following the final selection round, phage pools were analyzed by next-generation sequencing to identify enriched peptide sequences. Sequence analysis revealed recurrent CX_9_C-containing clones that increased in frequency over the course of selection, consistent with preferential enrichment of TREM2-binding peptides. Representative enriched clones were designated and prioritized for downstream synthesis and functional characterization.

To assess clone-level binding, selected phages were evaluated by phage ELISA against immobilized recombinant TREM2 ECD. Briefly, target-coated wells were incubated with individual phage clones, washed to remove unbound material, and bound phage were detected using an anti-M13 horseradish peroxidase-conjugated antibody and colorimetric substrate. Signal intensities were compared with background controls to identify clones exhibiting specific binding to TREM2. Clones showing reproducible enrichment and positive phage ELISA signals were advanced for chemical synthesis and orthogonal biophysical validation.

### Peptide synthesis and characterization

Peptide synthesis was carried out by solid phase peptide synthesis (SPPS) via the Fmoc/tBu strategy using an automated microwave peptide synthesizer - Liberty BLUE (CEM, Matthews, NC, USA). 424 mg Rink Amide ProTide Resin resin (0.59 mmol/g; CEM, Matthews, NC, USA) was used to synthesize each peptide. During the synthesis, each of the amino acids was coupled twice, using a 5-fold excess of the amount of resin deposition. The following solutions were used during the synthesis: OxymaPure/DIC as coupling reagents, 20% piperidine in DMF for Fmoc-deprotection, and DMF to wash between the deprotection and coupling steps. Synthesized peptides were cleaved from the resin using a mixture containing: 88% TFA, 5% H2O, 5% phenol and 2% TIPSI (10 ml of solution was used for 424 mg of resin). Crude peptides were precipitated with cold diethyl ether, decanted, and lyophilized. For purification, the peptides were dissolved in water with the addition of a 10-fold molar excess of dithiothreitol (DTT) over free sulfhydryl groups and incubated for 30 minutes at 60 °C. Peptides were purified by reversed phase HPLC, XBridge Prep C18 column (19 x 150 mm, 5 μm, Waters, MA, USA), flow rate 20 mL/min, 10 min run, 5-60% acetonitrile in water with 0.1% formic acid. The purity and mass spectra of the final products were analyzed using an ACQUITY HPLC system equipped with an SQ Detector 2 and an XBridge Prep C18 analytical column (4.6 × 150 mm, 5 μm, Waters, MA, USA), employing a linear gradient from 5% to 100% acetonitrile in water containing 0.1% formic acid over 10 minutes. Oxidation of the peptides was performed using compressed air. The peptide was dissolved in H2O and methanol (1:9, v:v), at a concentration of about 40 mg/L, and the pH was adjusted and kept between 8 and 9 using ammonia. The solution was stirred at room temperature for 7 days, and compressed air was run through the solution. After this time, the solvents were evaporated, and the peptides were lyophilized. Reaction progress was checked using analytical ACQUITY HPLC system equipped with an SQ Detector 2. After this process, the peptides were purified again using the same ACQUITY HPLC system on a XBridge Prep C18 column (19 x 150 mm, 5 μm, Waters, MA, USA). A linear gradient 5%-60% acetonitrile in water with 0.1% formic acid over 10 min was used.

### Monolith binding assay

#### Protein labeling

A 10 µM solution of the His-tagged TREM2 ectodomain (SinoBiological, 11084-H08H) was labeled using the RED-maleimide 2^nd^ generation dye (NanoTemper, MO-L014) according to the manufacturer’s protocol (labeling buffer: 10 mM HEPES pH 7.4, 150 mM NaCl, 0.5 mM TCEP, 0.05% Tween-20), resulting in a 0.90 µM stock of labeled protein (“TREM2-maleimide-dye”) in storage buffer (HBSP: 10 mM HEPES pH 7.4, 150 mM NaCl, 0.05% Tween-20) with a final degree of labeling (DOL) of 0.596. The labeled protein was aliquoted and stored at -80 °C until use.

#### Single-dose screening

TREM2-maleimide-dye was thawed and centrifuged for 10 min at 15,000 x g and 4 °C before use. The protein was diluted to 80 nM with assay buffer (HBSN: 10 mM HEPES pH 7.4, 150 mM NaCl). The dissolved peptides (10 mM in water) were diluted to 100 µM in assay buffer and mixed 1:1 with the protein solution (final concentrations: 40 nM TREM2-maleimide-dye, 50 µM peptide), followed by a 30-minute incubation at room temperature in the dark. Additionally, TREM2-maleimide-dye in assay buffer (HBSN) and TREM2-maleimide-dye in assay buffer with 2% DMSO were added as negative controls, and compound **T2337** at 50 µM in assay buffer with 2% DMSO was used as a positive control.^24^ The samples were transferred to Monolith Premium capillaries (NanoTemper, MO-K025) and measured at medium MST power and 100% excitation. The raw data were analyzed using GraphPad Prism 10. Peptides with a spectral shift (SpS) ratio (670 nm/650 nm) outside of a five standard deviation range from the average negative control (TREM2-maleimide-dye in HBSN, n = 4) were considered as potential hits. All TRIC traces were recorded for quality control (**Figure S1**). Samples that were flagged by the Monolith Control.2 software for aggregation or scan anomalies were excluded.

#### Control experiments

RED-maleimide 2^nd^ generation dye (NanoTemper, MO-L014) was diluted to 80 nM in assay buffer (HBSN pH 7.4). The dissolved peptides (10 mM in water) were diluted to 100 µM in assay buffer and mixed 1:1 with the dye solution (final concentrations: 40 nM RED-maleimide 2^nd^ generation dye, 50 µM peptide), followed by a 10-min incubation at room temperature in the dark. The samples were transferred to Monolith Premium capillaries and measured as described above. The raw data were analyzed using GraphPad Prism 10. Peptides outside of a 20% range from the initial fluorescence of the negative control (HBSN and dye, n = 4) were excluded due to assay interference. Peptides with an SpS ratio outside of a three standard deviation range from the negative control (HBSN and dye, n = 4) were excluded as well.

#### Binding affinity

TREM2-maleimide-dye was handled as described in the “Single-dose screening” section. The peptides were diluted to 200 µM in assay buffer (HBSN) and a subsequent 16-point dilution series was prepared. The samples were mixed 1:1 with the diluted protein and incubated for 30 minutes at room temperature in the dark before transferring them to Monolith Premium capillaries. All samples were measured at medium MST power and 100% excitation. The raw data were analyzed with GraphPad Prism 10 based on the instrument’s SpS (ratio 670 nm/650 nm) readout. All peptides were tested in three independent experiments.

### Cell-based evaluation of biological activity

#### Preparation of Amyloid-β Oligomers

Amyloid-β_1-42_ (Aβ_1-42_) peptide was dissolved in hexafluoro-2-propanol (HFIP), aliquoted, and dried under a gentle stream of nitrogen. The peptide film was resuspended in dimethyl sulfoxide (DMSO) and diluted in phenol-red-free cell culture medium to induce oligomer formation, followed by incubation at 4 °C for 24 h. Prepared Aβ oligomers were used immediately for cellular assays.

#### Human iPSC-Derived Microglia Culture and Treatment

Human induced pluripotent stem cell (iPSC)-derived microglia were cultured according to the supplier’s protocol in microglia maintenance medium. Cells were seeded into 96-well plates and allowed to equilibrate for 24 h prior to treatment. The tested peptides were prepared as PBS stock solutions and diluted into culture medium to final concentrations of 5, 10, and 25 µM (n=5). Microglia were pretreated with compounds for 1 h, followed by stimulation with Aβ_1-42_ oligomers. After 24 h, culture supernatants were collected for cytokine analysis. Vehicle-treated cells with and without Aβ stimulation served as controls. WT and KO microglia were treated identically in all experiments.

#### Measurement of IL-1β Secretion

Interleukin-1β (IL-1β) levels in microglial culture supernatants were quantified using a commercially available human IL-1β ELISA kit (from Abcam, Catalog# ab214025) according to the manufacturer’s instructions. Absorbance was measured using a microplate reader, and cytokine concentrations were calculated from a standard curve. Data were normalized to the Aβ-stimulated vehicle control and expressed as percent change relative to this condition.

#### Measurement of Secreted ApoE

ApoE levels in microglial culture supernatants were quantified using a commercially available human ApoE ELISA kit (from Abcam, Catalog # ab108813) according to the manufacturer’s instructions. Supernatants were collected 24 h after Aβ stimulation and compound treatment.

ApoE concentrations were calculated from a standard curve and normalized to Aβ-treated vehicle controls.

#### Neuron-Microglia Co-culture Assay

Human iPSC-derived neurons were cultured on poly-D-lysine/laminin-coated plates until mature neuronal networks were established. iPSC-derived microglia were added to neuronal cultures at a neuron-to-microglia ratio of 5:1 and allowed to integrate for 24 h. Co-cultures were treated with the tested peptides (5, 10, or 25 µM, n=5) or positive control (VG3927, 5 μM), followed by exposure to Aβ_1-42_ oligomers for 48-72 h.

#### Quantification of PSD95 Levels

PSD95 levels were quantified using Human PSD95 ELISA Kit from Abcam (Catalog No. 50-269-8625). Data were expressed as percent rescue relative to Aβ-treated vehicle controls and analyzed using GraphPad Prism.

#### Statistical Analysis

All experiments were performed with a minimum of three independent replicates. Data are presented as mean ± standard deviation (SD). Statistical comparisons between groups were performed using one-way analysis of variance (ANOVA) followed by Dunnett’s post hoc test. Differences were considered statistically significant at *p* < 0.05.

## Computational studies

Initial structures of TREM2 in complex with cyclic peptides **TREM2-6** (**T6**) and **TREM2-12** (**T12**) were generated using AlphaFold3.^28^ Molecular dynamics (MD) simulations were then run to test the validity of the predictions and assess the dynamic stability of each of the TREM2-p6 and TREM2-**T12** receptor-peptide complexes at 300K for 100ns. Each system consisted of the TREM2 receptor (TREM2-apo) alone, or as part of one of two complexes (TREM2-**T6**, TREM2-**T12**). All simulations were run by combining the AMBER software package^29,30^ with the ff14SB force field^31^ for protein and peptide parameters. Water TIP3P^32^ was used as solvent, and all solutes were contained within a truncated octahedral box that had been cut off 15.0Å from the solute surface. Ions (sodium and chloride) were also included to neutralise the charge of the systems.

Before running the full MD simulations, each of the systems underwent an energy minimisation process to reduce unfavourable contacts in the system. This process consists of two stages. First, 5000 steps of steepest descent (SD) followed by 5000 steps of conjugate gradient (CG) energy optimisations were performed with position constraints applied to the solutes. Second, all constraints were released, and the second stage of the energy minimisation was completed, with another 5000 steps of SD and 5000 steps of CG.

Following the energy minimisations, the systems were heated from 0K to 300K in 100ps. Afterwards, the systems were subjected to a 100 ns production run, under constant number of particles, temperature, and pressure conditions (NPT ensemble),^33^ at 300K and 1.0bar. Temperature control was achieved through the use of a Langevin thermostat^34^ (collision frequency of 2.0ps-1), while pressure regulation was adjusted accordingly. Hydrogen bonds were constrained using the SHAKE algorithm,^35^ thereby allowing a 2fs time step. Trajectory analysis, including RMSD and hydrogen bonding maps, was accomplished using the CPPTRAJ module of AmberTools.

## Evaluation of PK and physicochemical properties

These experiments were conducted following our previously reported methods.^22^

## AUTHOR INFORMATION

### Corresponding Author

Moustafa T. Gabr - Department of Radiology, Molecular Imaging Innovations Institute (MI3), Weill Cornell Medicine, New York, NY 10065, United States;

## ASSOCIATED CONTENT

The following files are available free of charge.

Raw data for MST studies, control experiments for TREM2 binding, individual peptide binding curves, and chemical characterization of the synthesized compounds (PDF).

## Notes

The authors declare no competing financial interest.

