## Supporting Information for "Phage Display-Derived Cyclic Peptides Target TREM2 and Modulate Microglial Responses under Amyloid Stress"

*Electronic Supplementary Material for*

Chemical characterization of the synthesized peptides……………………………………………………….…S4

### Monolith binding assay

#### Single-dose screening

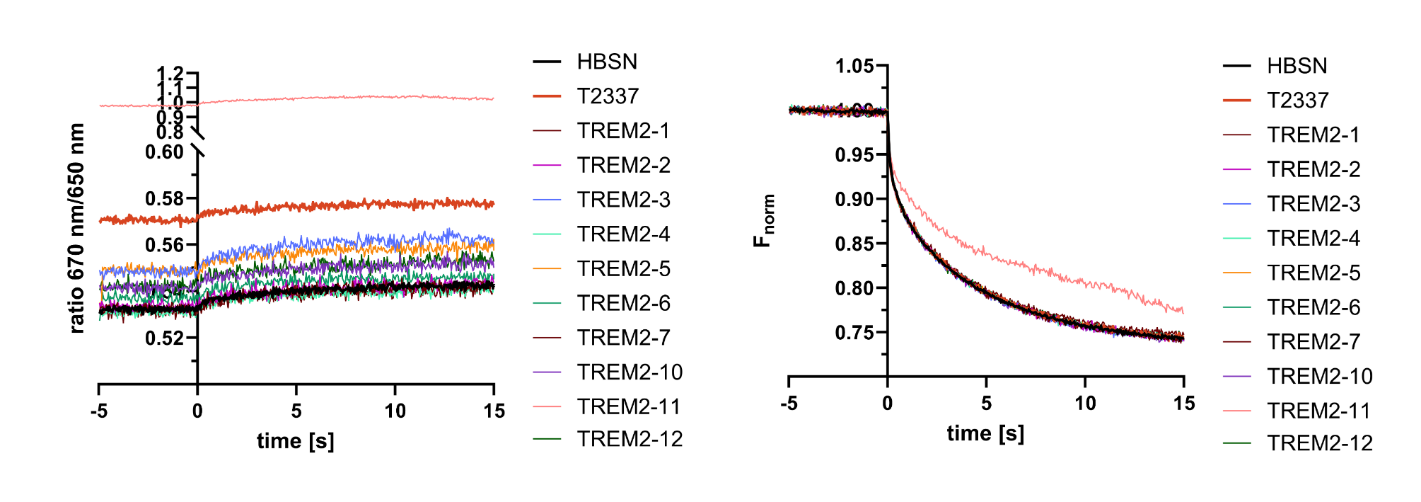

**Figure S1.** **Raw data for single-dose screening.** Left: SpS traces for all controls (negative: HBSN, positive: **T2337**) and peptides. Right: TRIC traces for quality control, visible aggregation for peptide **TREM2-11**. All graphs were created with GraphPad Prism 10.

#### Control experiments

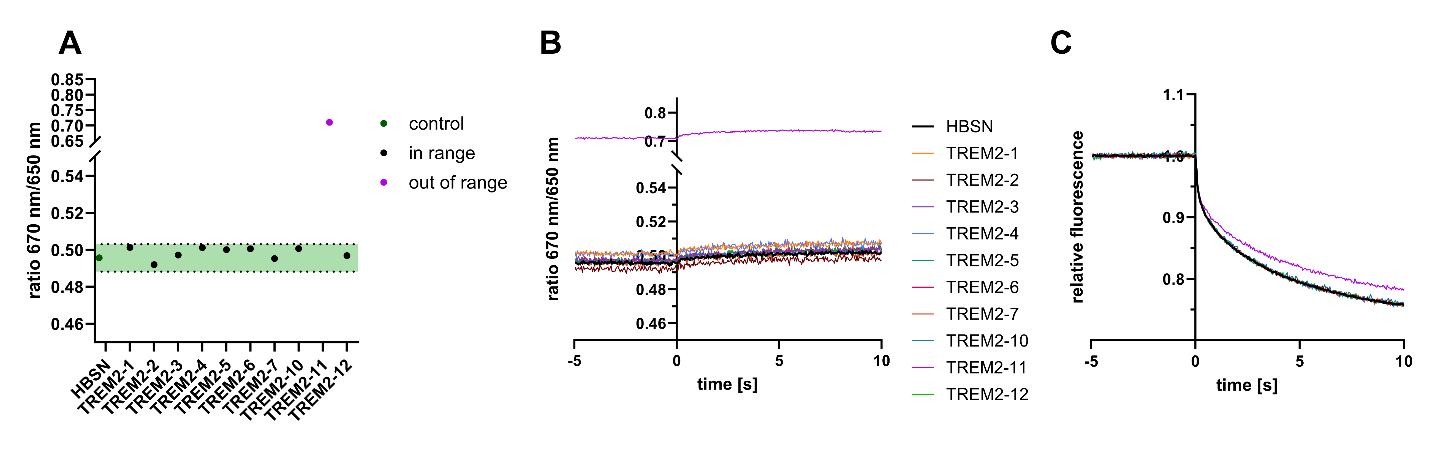

**Figure S2.** **Results of control experiments with RED-maleimide-dye (2^nd^ generation).** (**A**) Spectral shift plot for average control (HBSN, n = 4) plus five standard deviation range shaded in green. Samples outside of range are highlighted in purple. (**B**) Raw SpS traces for the control experiments with RED-maleimide-dye. (**C**) Raw TRIC traces for the control experiments with RED-maleimide-dye. All graphs plotted with GraphPad Prism 10. Figure composed using BioRender.

#### Binding affinity studies

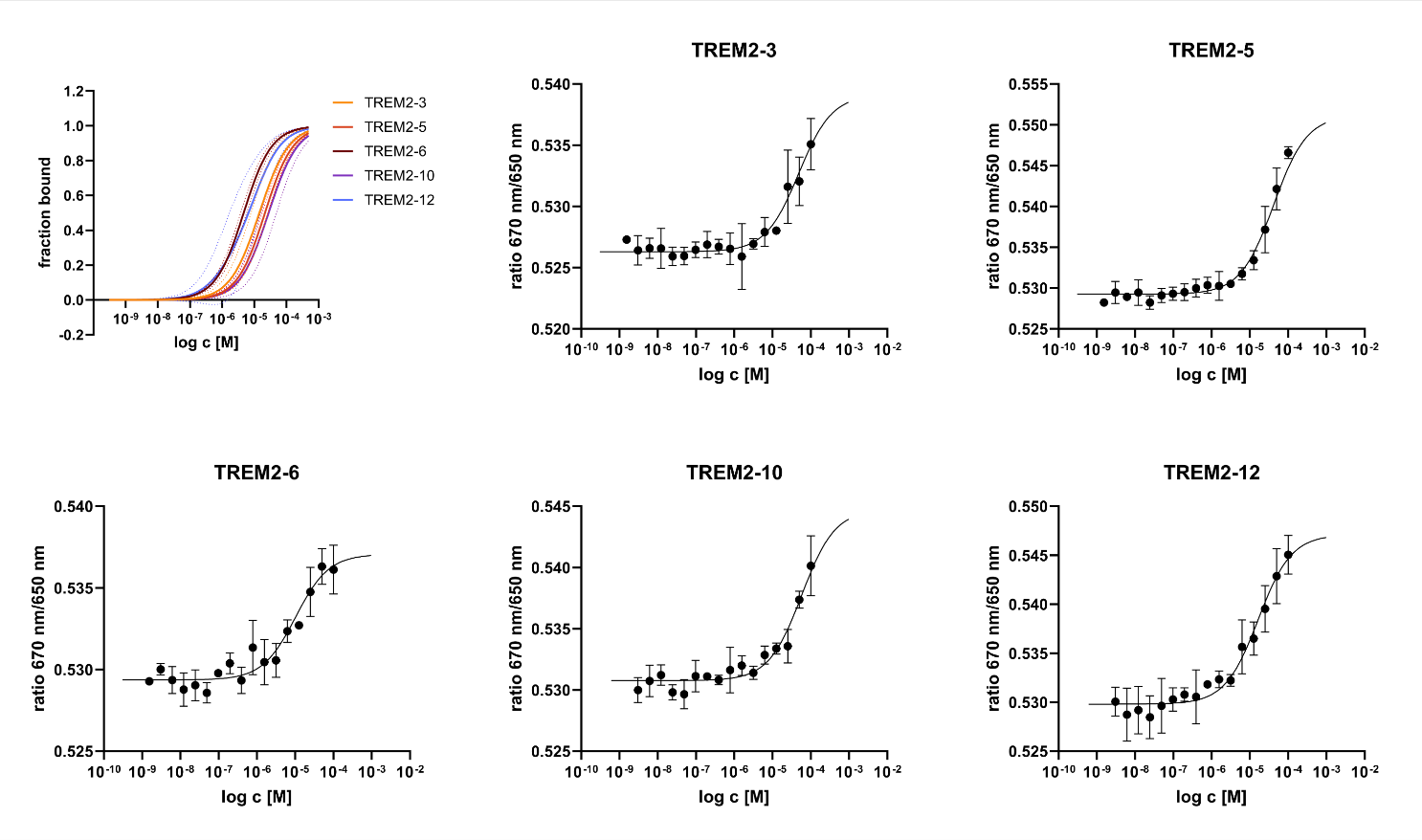

**Figure S3.** Individual graphs for all five hit peptides displayed as spectral shift (SpS, ratio 670 nm/650 nm) vs. log c [M]. The graphs were normalized to fraction bound vs. log c [M] for better comparability (top left). All graphs were plotted with GraphPad Prism 10. Figure composed using BioRender.

**TREM2-1 LC**

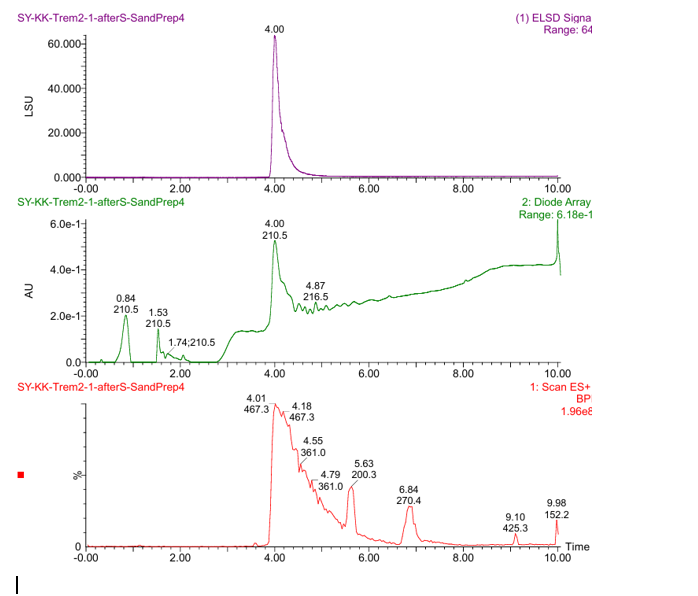

**TREM2-1 MS**

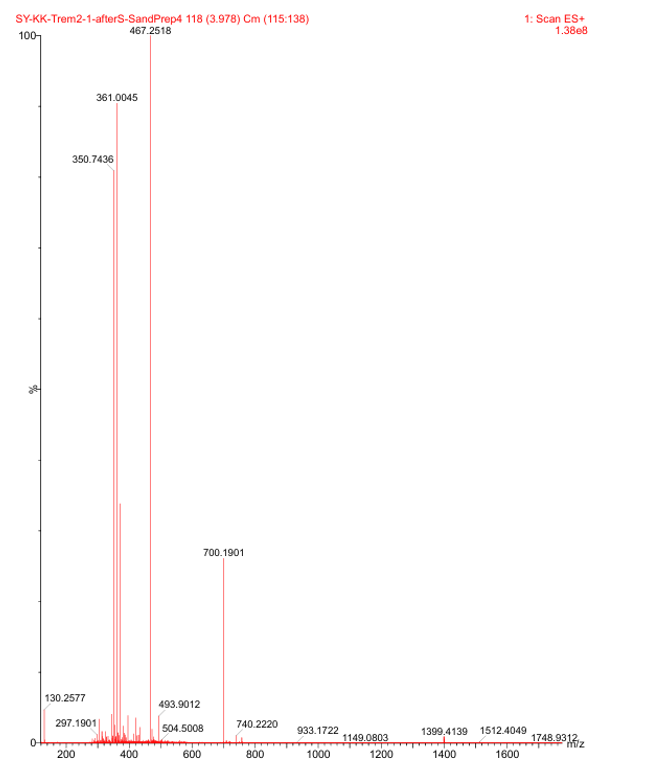

**TREM2-3 LC**

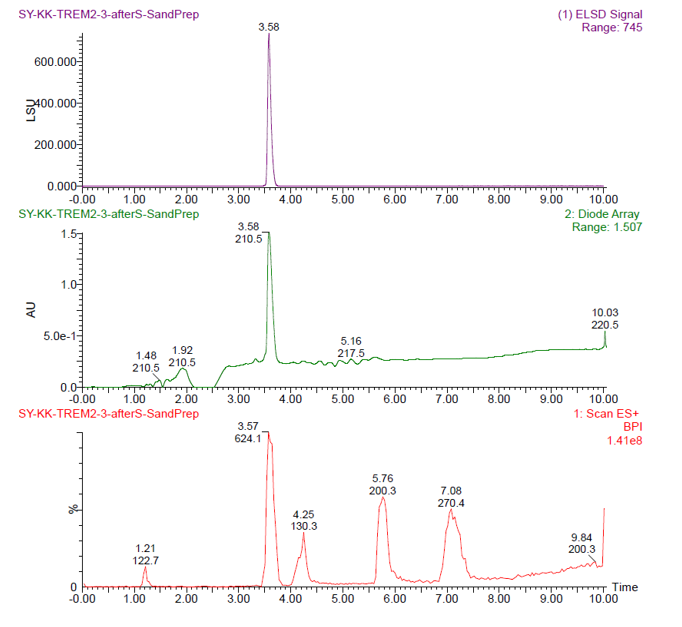

**TREM2-3 MS**

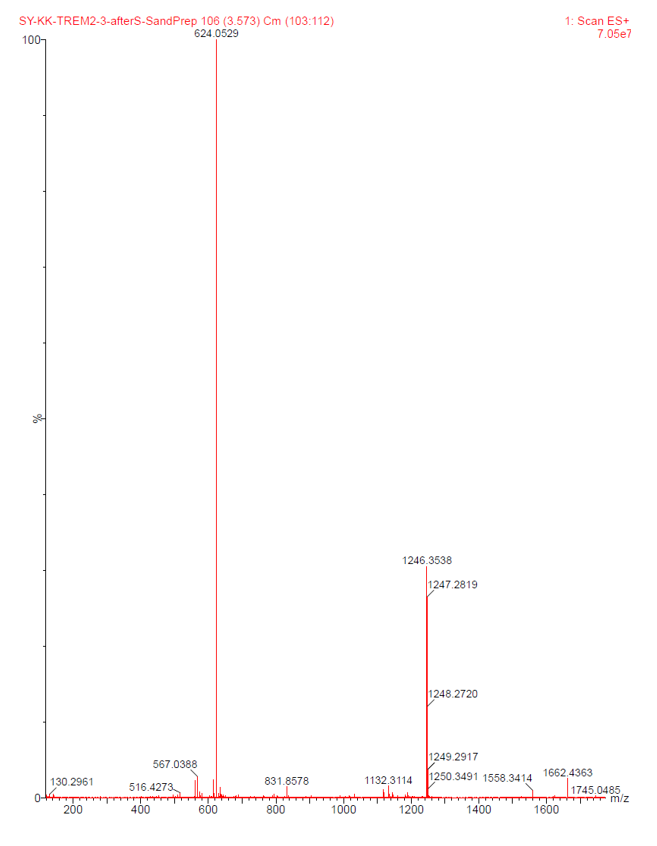

**TREM2-4 LC**

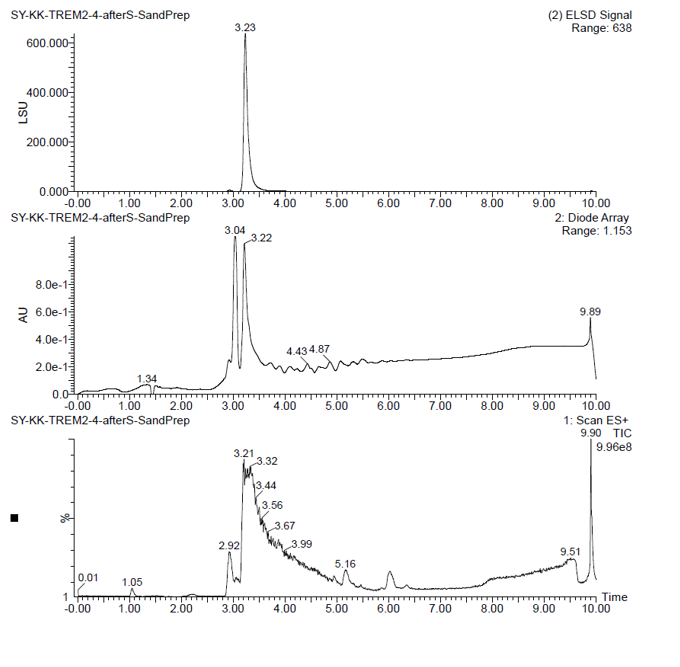

**TREM2-4 MS**

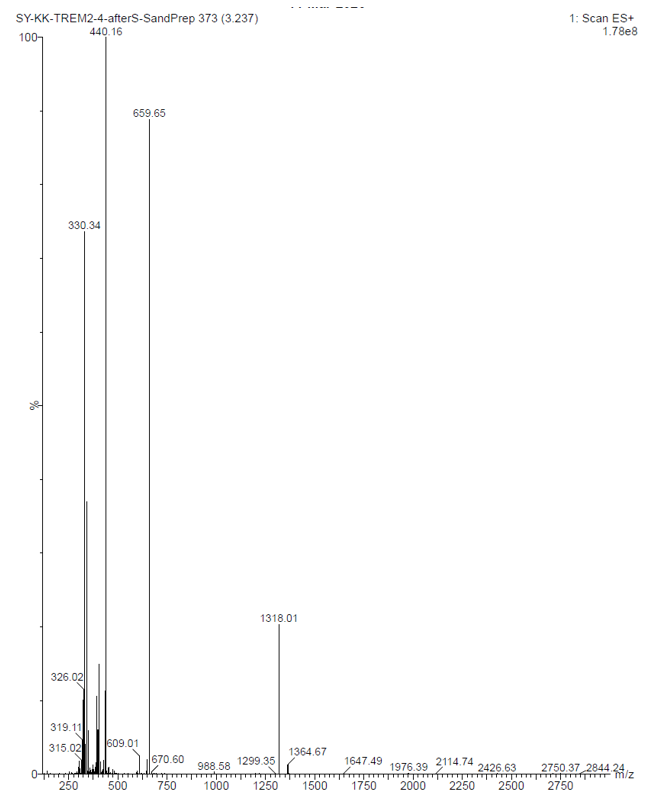

**TREM2-5 LC**

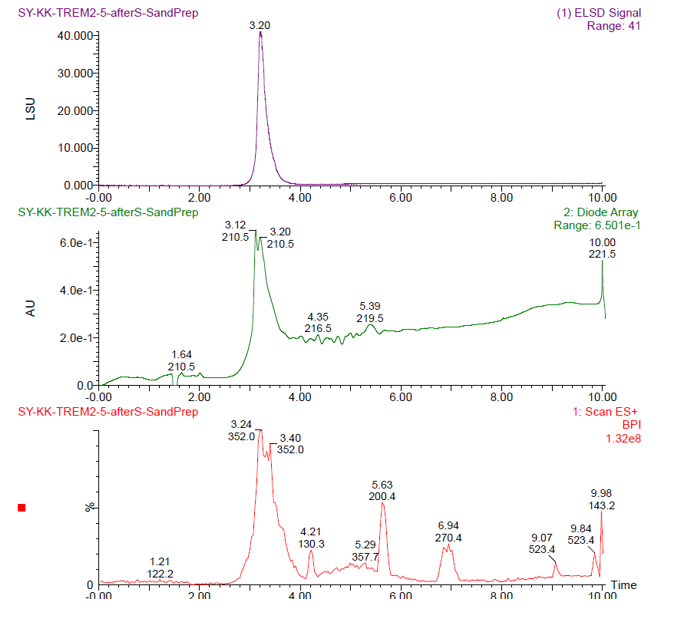

**TREM2-5 MS**

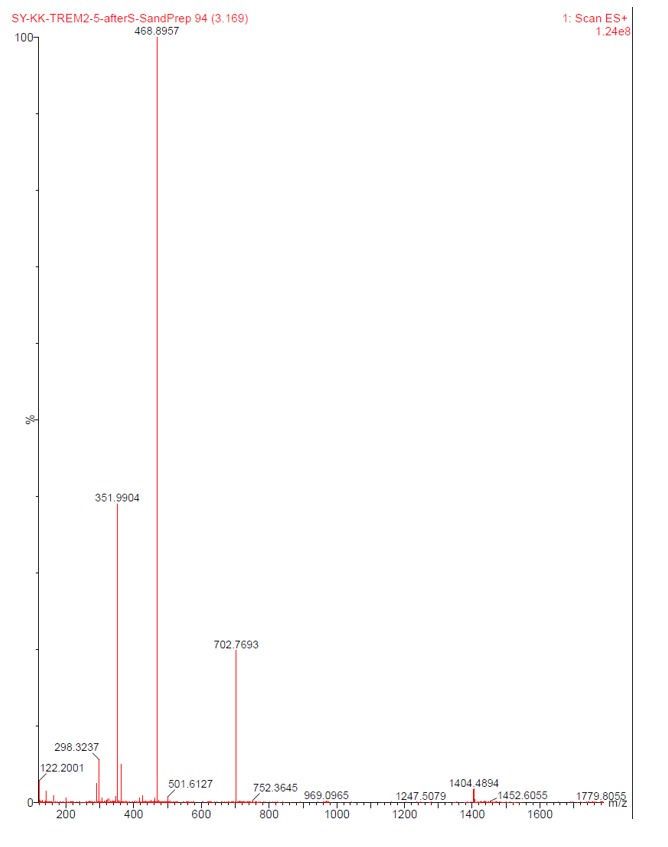

**TREM2-6 LC**

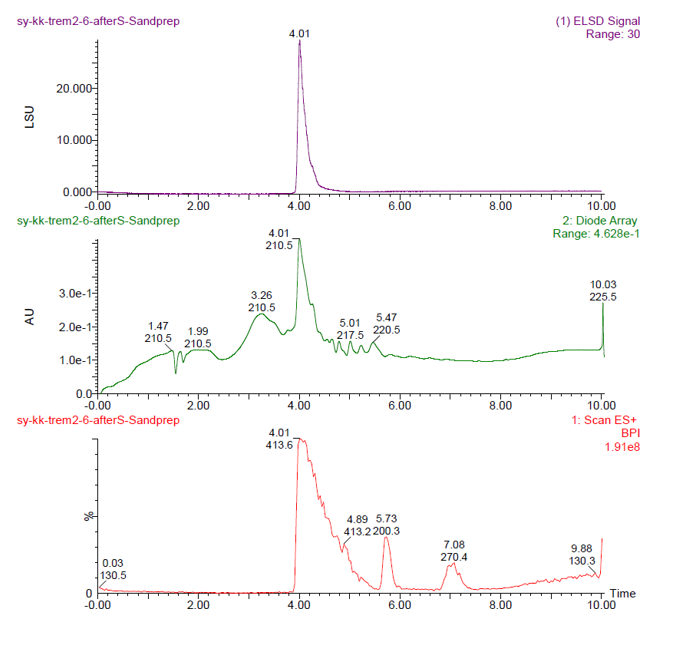

**TREM2-6 MS**

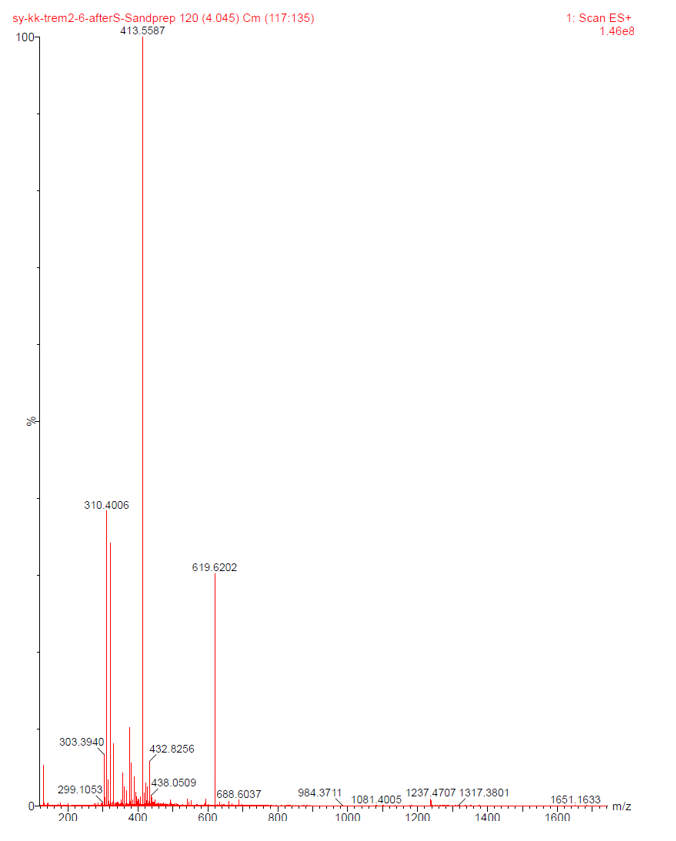

**TREM2-7 LC**

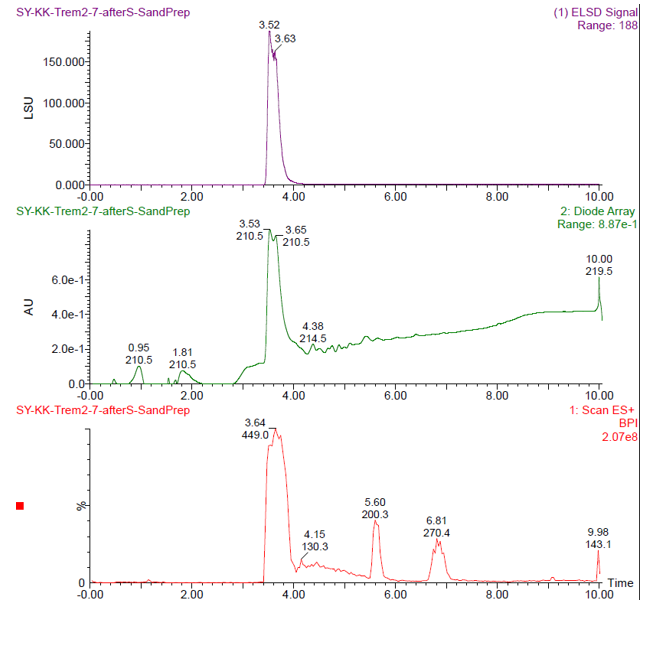

**TREM2-7 MS**

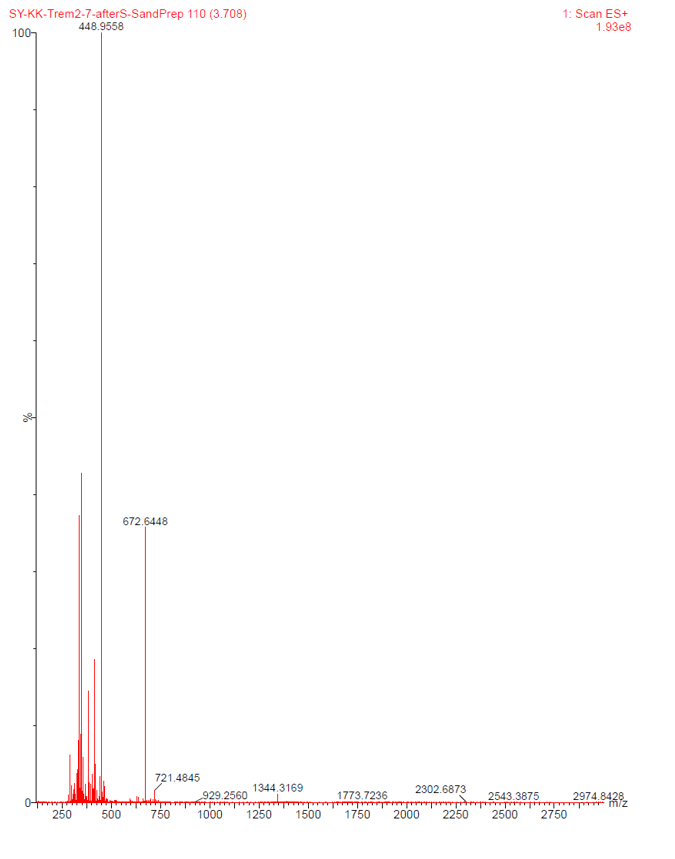

**TREM2-10 LC**

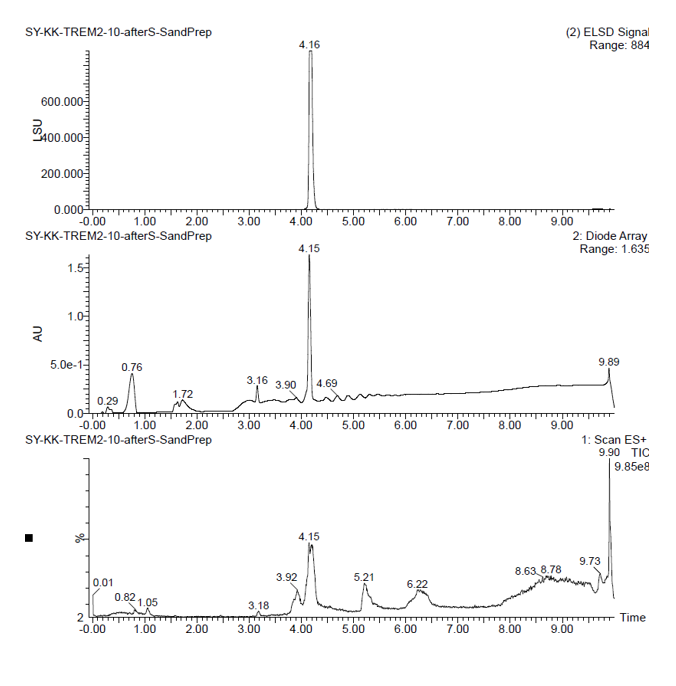

**TREM2-10 MS**

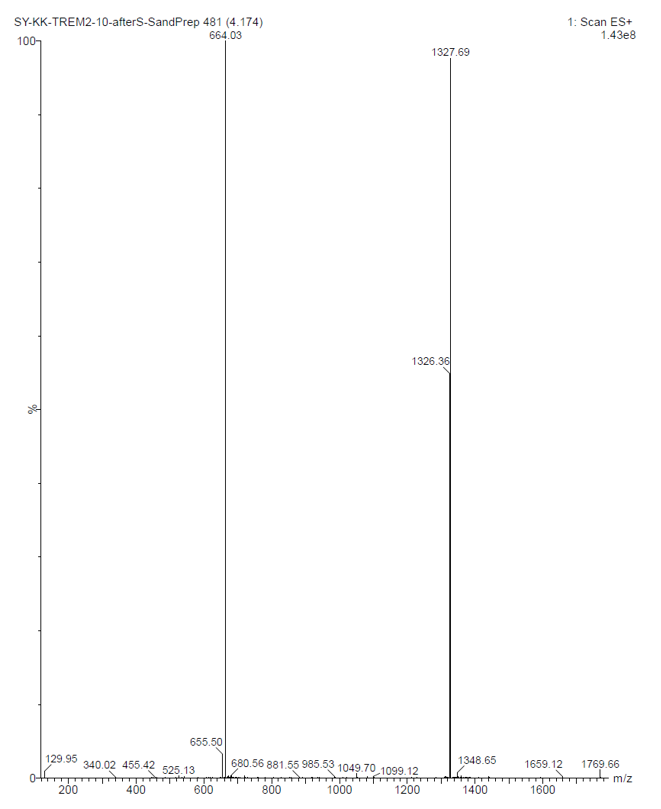

**TREM2-11 LC**

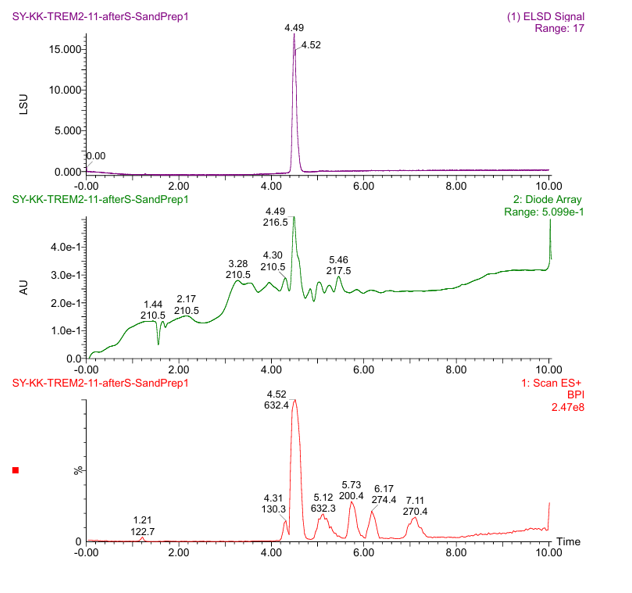

**TREM2-11 MS**

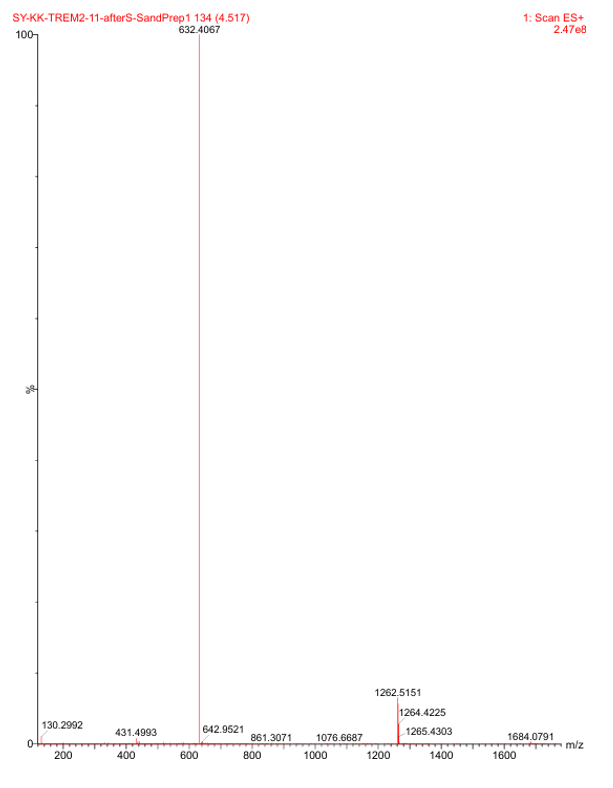

**TREM2-12 LC**

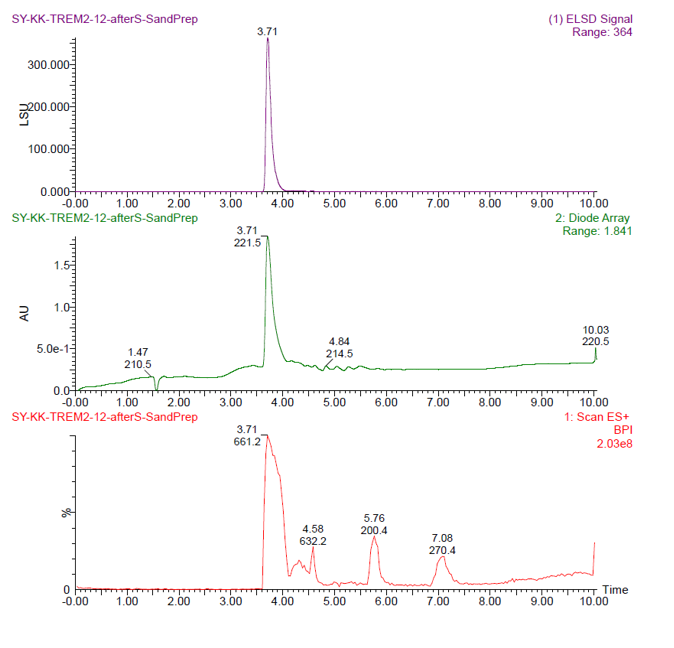

**TREM2-12 MS**

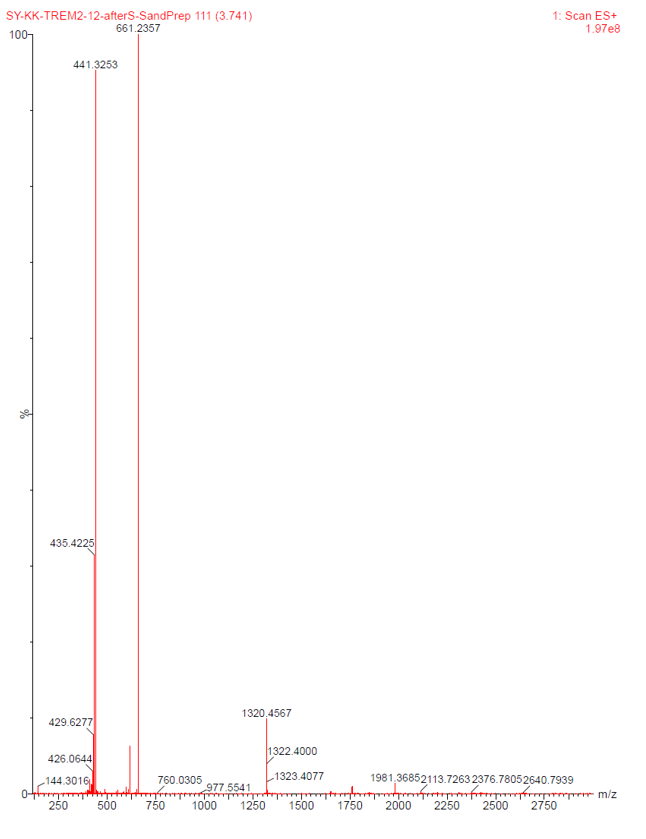
